# Comparative reservoir competence of *Peromyscus leucopus*, C57BL/6J, and C3H/HeN for *Borrelia burgdorferi* B31

**DOI:** 10.1101/2023.09.28.559638

**Authors:** Jeffrey S. Bourgeois, Stephanie S. You, Luke H. Clendenen, Muskan Shrestha, Tanja Petnicki-Ocwieja, Sam R Telford, Linden T. Hu

## Abstract

*Borrelia burgdorferi*, a Lyme disease spirochete, causes a range of acute and chronic maladies in humans. However, a primary vertebrate reservoir in the United States, the white-footed deermouse *Peromyscus leucopus*, is reported not to have reduced fitness following infection. While laboratory strains of *Mus musculus* mice have successfully been leveraged to model acute human Lyme disease, the ability for these rodents to model *B. burgdorferi*-*P. leucopus* interactions remains understudied. Here we compared infection of *P. leucopus* with *B. burgdorferi* B31 with infection of the traditional *B. burgdorferi* murine models—C57BL/6J and C3H/HeN *Mus musculus*, which develop signs of inflammation akin to human disease. We find that *B. burgdorferi* were able to reach much higher burdens (10- to 30-times higher) in multiple *M. musculus* skin sites, and that the overall dynamics of infection differed between the two rodent species. We also found that *P. leucopus* remained transmissive to larval *Ixodes scapularis* for a far shorter period than either *M. musculus* strain. In line with these observations, we found that *P. leucopus* does launch a modest but sustained inflammatory response against *B. burgdorferi* in the skin, which we hypothesize leads to reduced bacterial viability and rodent-to-tick transmission in these hosts. Similarly, we also observe evidence of inflammation in infected *P. leucopus* hearts. These observations provide new insight into reservoir species and the *B. burgdorferi* enzootic cycle.

**Importance:** A Lyme disease-causing bacteria, *Borrelia burgdorferi*, must alternate between infecting a vertebrate host—usually rodents or birds—and ticks. In order to be successful in that endeavor the bacteria must avoid being killed by the vertebrate host before it can infect a new larval tick. In this work we examine how *B. burgdorferi* and one of its primary vertebrate reservoirs, *Peromyscus leucopus*, interact during an experimental infection. We find that *B. burgdorferi* appear to colonize its natural host less successfully than conventional laboratory mouse models which aligns with a sustained seemingly anti-bacterial response by *P. leucopus* against the microbe. These data enhance our understanding of *P. leucopus* host-pathogen interactions and could potentially serve as a foundation to uncover ways to disrupt the spread of *B. burgdorferi* in nature.

## Introduction

Few endemic pathogens in the United States are as infamous as *Borrelia burgdorferi* sensu stricto (also called *Borreliella burgdorferi*), a Lyme disease spirochete which causes a range of acute and late disease symptoms in humans. In North America, *B. burgdorferi* is spread between vertebrate hosts by the deer tick or black-legged tick *Ixodes scapularis* (1-3). This enzootic cycle occurs ad infinitum as larval ticks acquire *B. burgdorferi* from infected vertebrates, molt, transmit the infection to new vertebrate hosts as nymphal ticks, molt into adult ticks, and produce new uninfected larval ticks (4). While *B. burgdorferi* can infect numerous permissive vertebrate hosts ranging from birds (5) to dogs (6), one of the dominant North American reservoir hosts for *B. burgdorferi* is the white-footed deermouse *Peromyscus leucopus* (7). By some estimates, *P. leucopus* accounts for >30% of *B. burgdorferi* acquired by larval ticks (8), though the exact value likely varies by region and year. This aligns with other work which noted that in endemic regions, up to 30-95% of this highly abundant rodent are infected with *B. burgdorferi* (9, 10). Numerous ecological and laboratory studies have noted a unique relationship between *B. burgdorferi* and *P. leucopus* as, outside of neonate infection (11), *B. burgdorferi* has no detectable impact on *P. leucopus* fitness (10, 12) and has not been reported to drive any observed behavioral (13, 14) or histological (11) changes. Even at the molecular level, two studies found very few changes to gene expression during post-acute infection (4-5 weeks post infection) in skin (15) or late infection (10 weeks post infection) in spleen (16). The transcriptomic inflammatory response is limited despite considerable evidence that *P. leucopus* remain colonized by *B. burgdorferi* for months following infection (10, 17, 18).

In the laboratory, human Lyme disease is often modeled using inbred *Mus musculus* mouse models in which infection of some strains (*e*.*g*. C3H mice) results in substantial carditis and arthritis, while other strains (C57BL/6) display more modest inflammation (19, 20). However, the capacity for *M. musculus* to model the enzootic cycle is less well understood. Work by Hanincova *et al*. suggested that *P. leucopus* are not as effective at infecting ticks with *B. burgdorferi* compared with C3H laboratory mice (18). However, longitudinal studies quantifying bacterial burden, host inflammation, and the ability to transmit from rodents to ticks across these two species have not been performed. Here we seek to examine the relationship between *P. leucopus* and *B. burgdorferi* by examining host and bacterial dynamics throughout infection and compare the interactions to those of more commonly used laboratory mice. Using the *B. burgdorferi* B31 type-strain, we assess *B. burgdorferi* burden dynamics between laboratory-reared *P. leucopus* and two inbred *Mus musculus* mouse strains (C57BL/6J and C3H/HeN). We find that *P. leucopus* have lower *B. burgdorferi* burden in ear tissue compared to the laboratory mouse strains. This correlates with a modest transcriptional burst of genes associated with innate and adaptive immunity in the skin and very mild carditis in *P. leucopus*, suggesting that *P. leucopus* are actively responding to and controlling *B. burgdorferi* B31 infection. Notably, infected *P. leucopus* transmit less *B. burgdorferi* to *I. scapularis* larvae than similarly infected laboratory mouse strains. Together, these data demonstrate that despite being an important reservoir host for *B. burgdorferi, P. leucopus* exert pressure on *B. burgdorferi* B31 during laboratory tick-to-rodent-to-tick cycling.

## Results

### *Peromyscus leucopus* restrict *Borrelia burgdorferi* B31 during early infection

Work in other laboratories has demonstrated that *P. leucopus* respond to the bacterial pathogen associated molecular pattern lipopolysaccharide (LPS) to a far lesser extent than inbred (BALB/c) (21) and outbred (CD-1) (22) *M. musculus*. While *B. burgdorferi* lacks LPS, if bacterial sensing is broadly suppressed in *P. leucopus, B. burgdorferi* could be allowed to thrive until the bacteria is collected during another tick bloodmeal. To test whether *P. leucopus* is more permissive to *B. burgdorferi* during early infection, we injected laboratory-reared *P. leucopus* and two inbred *M. musculus* strains (C57BL/6J and C3H/HeN) subcutaneously with 1×10^5^ *B. burgdorferi* B31 spirochetes and collected skin from the injection site one week later. Note that these *M. musculus* strains were chosen because they have routinely been used to model *B. burgdorferi* infections (23-28), and C3H have previously been used as a comparator when studying *Peromyscus*-*Borrelia* interactions (16, 18, 29). *B. burgdorferi* strain B31was chosen due to its place as the most commonly used wild-type strain which allows comparisons to other studies.

We were able to culture spirochetes from the injection site of all the tested rodents **(Figure 1A)**, however we found that *P. leucopus* had equal bacterial burdens (calculated by quantifying bacterial genomic DNA copies by digital droplet PCR [ddPCR]) at the injection site as C57BL/6J mice. C3H/HeN mice had higher, albeit more variable, burden compared to *P. leucopus* **(Figure 1B)**.

**Figure 1:**
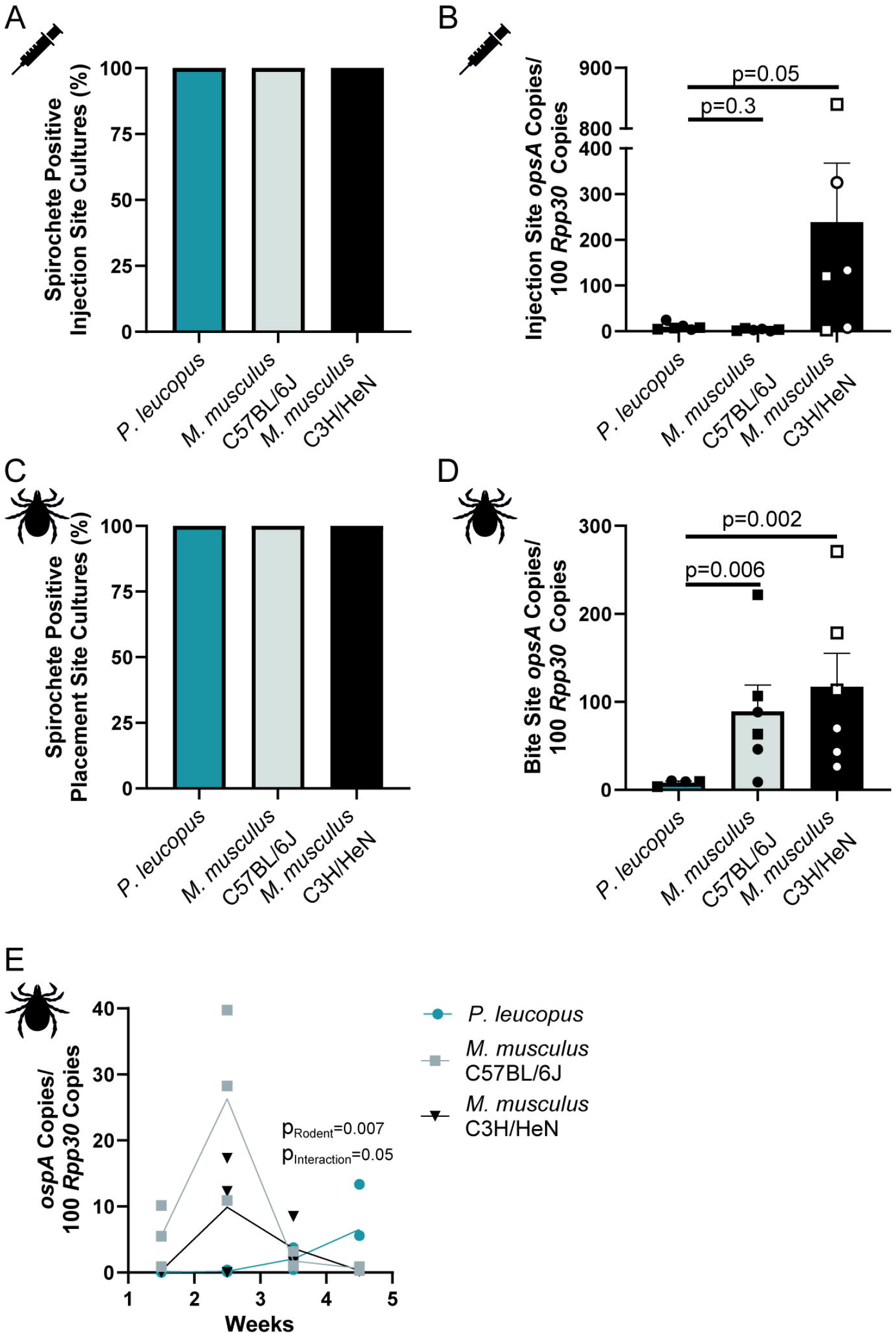
*B. burgdorferi* B31 expand to greater numbers in *M. musculus* than in *P. leucopus* following tick bite. (A) *B. burgdorferi* can be cultured from needle injection sites from *P. leucopus* and *M. musculus* (n=6) one week post injection. (B) *B. burgdorferi* are present at substantially lower numbers at the needle injection site in *P. leucopus* than C3H/HeN mice by ddPCR. For panels A and B rodents were injected with 100,000 *B. burgdorferi* via subcutaneous injection. (C) *B. burgdorferi* can be cultured from tick placement sites one week post infestation. (D) *B. burgdorferi* are present at substantially lower numbers at the tick placement site in *P. leucopus* (n=4) than C57BL/6J (n=6) or C3H/HeN (n=6) one week post infestation by ddPCR. For panels C and D, infected nymphal *I. scapularis* were used to infect rodents. The tick placement region was harvested one week post infestation. For panels A and C, cultures were monitored for spirochetes for three weeks. For Panels B and D, *B. burgdorferi ospA* DNA copies are normalized to copies of the mammalian *Rpp30* gene and p-values are generated by one-way ANOVA on log transformed data with Dunnett’s multiple comparison test. For panels A-D, all rodents were between 8-12 weeks old at time of infection and both sexes were present. For panels A and D, bars represent the mean and error bars the standard error of the mean. Individual rodents are plotted as dots (circle = female, square = male). (E) *B. burgdorferi* has different population dynamics in ear tissue from male *P. leucopus* or *M. musculus* (n=3) following tick bite. Nymphal *I. scapularis* were used to infect rodents and ear punches were taken weekly starting at 1.5 weeks post tick placement for ddPCR analysis. P-value generated by two-way ANOVA on log transformed data. Individual rodents were plotted as dots and lines intercept the mean. For all panels, data are pooled from two independent experiments.

While the *B. burgdorferi* subcutaneous injection model offers many advantages, it does not always accurately recapitulate wild infection where *I. scapularis* deliver a small number of *B. burgdorferi* and saliva proteins into the host during the bloodmeal. This is particularly true in our experimental design above where we used a large dose to infect the rodents. In order to better replicate natural infections, we infected rodents by placing five infected *I. scapularis* nymphs on the backs of *P. leucopus* and *M. musculus*. We then harvested skin from the tick placement site one week later. Again, we successfully cultured spirochetes from all of the tick placement sites **(Figure 1C)**, but this time we found that *P. leucopus* had substantially (>10-fold) lower bacterial burdens at the placement site compared to either C57BL/6J or C3H/HeN mice **(Figure 1D)**. Thus, we conclude that despite being an important reservoir host for *B. burgdorferi* and *I. scapularis* in North America (7), our laboratory-reared *P. leucopus* are far less permissive to high *B. burgdorferi* B31 burdens immediately following tick bite.

To further characterize *B. burgdorferi* burdens in rodent skin, we tracked bacterial burdens in individual rodents infected by nymphal ticks over time, taking weekly ear punches over one month. We found that similar to our findings at the tick placement site, *P. leucopus* had reduced bacterial burdens at the disseminated ear site at early time points compared to the *M. musculus* strains **(Figure 1E)**. While both *M. musculus* strains had their peak infection at approximately 2.5 weeks post-nymph placement, we did not observe substantial increases in bacterial burdens in *P. leucopus* until 3.5-4.5 weeks post-nymph placement. This demonstrated that *P. leucopus* also appears to be less permissive to *B. burgdorferi* B31 in ear during the disseminated phases of infection and that the overall infection dynamics in *P. leucopus* are substantially different than in the *M. musculus* model as indicated by a significant interaction term in our Two-Way ANOVA (p_rodent x time_= 0.05).

### Examining *B. burgdorferi* and *P. leucopus* interactions over time

To better understand how *P. leucopus* and *B. burgdorferi* B31 interact with each other over the course of infection, we infested 20 *P. leucopus* with infected nymphs and 4 *P. leucopus* with uninfected nymphs across two independent experiments. Each rodent was euthanized at one of five terminal time points (1-, 2-, 4-, 8-, or 13-weeks post infestation, two rodents euthanized per time point per experiment, two independent experiments) to obtain ear and heart tissue for microbiological, genomic, transcriptomic, and histopathological analysis **(Figure 2A)**. Rodents infested with uninfected nymphs were harvested one week post placement. Additionally, each infected rodent was subjected to one intermediate time point (1-, 2-, 4-, 10-weeks post infestation) in which an ear punch was taken from a living rodent prior to infestation with larval *I. scapularis*. These time points were chosen because *P. leucopus* often become infected during the spring *I. scapularis* nymphal peak and transmit to larval *I. scapularis* 1-3 months later during the summer larval peak (30-32), and thus this captures the most important period of time that *B. burgdorferi* must persist in *P. leucopus* during the enzootic cycle. For comparison, we repeated these time courses with C57BL/6J and C3H/HeN *M. musculus*. Due to logistical constraints, these time courses were completed separately and thus caution should be used when directly comparing the *P. leucopus* and *M. musculus* data.

**Figure 2:**
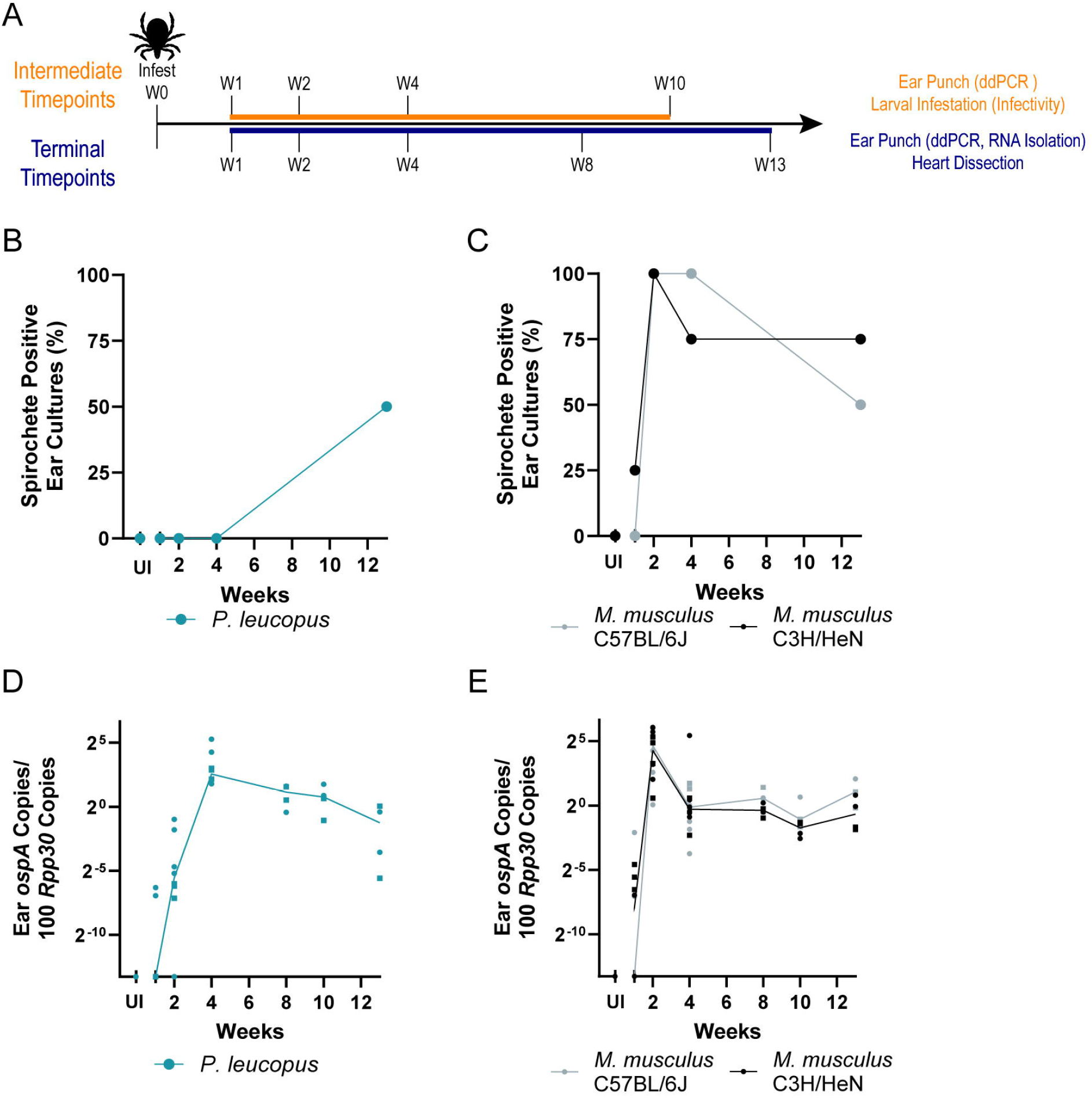
Population dynamics of *B. burgdorferi* B31 in *P. leucopus* and *M. musculus* over thirteen weeks. (A) Schematic of the experimental design used for Figures 2-5. Rodents were infested with five nymphal ticks (infected or uninfected) and rodents were harvested at the listed terminal time points. Rodents were subjected to a maximum of one intermediate time point. All rodents were between 6 and 12 weeks old at the time of infection. Each time point included one male and one female rodent per species per experiment where possible. All time points represent between three and eight samples depending on whether the time point combines terminal and intermediate values (*e*.*g*. week 2 n=8, week 8 n=4) and/or whether samples needed to be excluded for technical reasons (*e*.*g*., rodent death, failed DNA extraction). (B, C) *B. burgdorferi* can be cultured from *P. leucopus* ear punches late in infection (B), while *M. musculus* can be cultured from ear punches starting at two weeks post nymph placement (C). (D, E) The population dynamics of *B. burgdorferi* in ear tissue of *P. leucopus* (D) and *M. musculus* (E) ears are measured by ddPCR. The *B. burgdorferi ospA* gene copies are normalized to copies of the mammalian *Rpp30* gene. Each dot represents a single mouse (circles = female, squares = male), the connecting line intercepts the median value of each timepoint. Samples that did not have any copies of *ospA* are plotted at 0.0001 copies per 100 *Rpp30* copies.

At each terminal time point, we attempted to culture spirochetes from approximately one half of a rodent ear in BSK-II. Surprisingly, we were unable to culture *B. burgdorferi* B31 from *P. leucopus* until late in the infection **(Figure 2B)** compared to *M. musculus* **(Figure 2C)**. Quantification of bacterial burdens corroborated our previous observations **(Figure 1E)** in that *P. leucopus* had a modest spike in bacterial burden at four weeks post infestation **(Figure 2D)** while both lab mouse strains had substantially peaks at two weeks post infestation **(Figure 2E)**. Both species remain infected throughout the timecourse.

### *B. burgdorferi* rodent-to-larval tick transmission is lower in *P. leucopus* than *M. musculus*

To examine how the differences in skin burden over time correlated with the ability for each rodent to transmit *B. burgdorferi* to larval *I. scapularis*, we infested rodents with larval ticks at intermediate timepoints over our time course **(Figure 2A)**. Notably, we also collected ear punch biopsies from the rodents in order to calculate bacterial burden for each specific rodent at the time of infestation.

*P. leucopus* can transmit *B. burgdorferi* to larval ticks very soon after placement of infected nymphal ticks with ∼25% of larval ticks placed one week after infected nymph feeding yielding viable *B. burgdorferi* B31 **(Figure 3A)**. This increased to 80% at two weeks post nymph placement before sequentially dropping for the duration of the experiment (two-week vs ten-week timepoint p=0.0002). Notably there was no correlation between the observed bacterial burden in the ear by ddPCR and the proportion of infected larvae recovered from the rodents **(Figure 3B)**. Together these data suggest that in addition to having smaller numbers of viable *B. burgdorferi* B31 in the ear, *P. leucopus* may have additional mechanisms that inhibit the transmission of *B. burgdorferi* to ticks.

**Figure 3:**
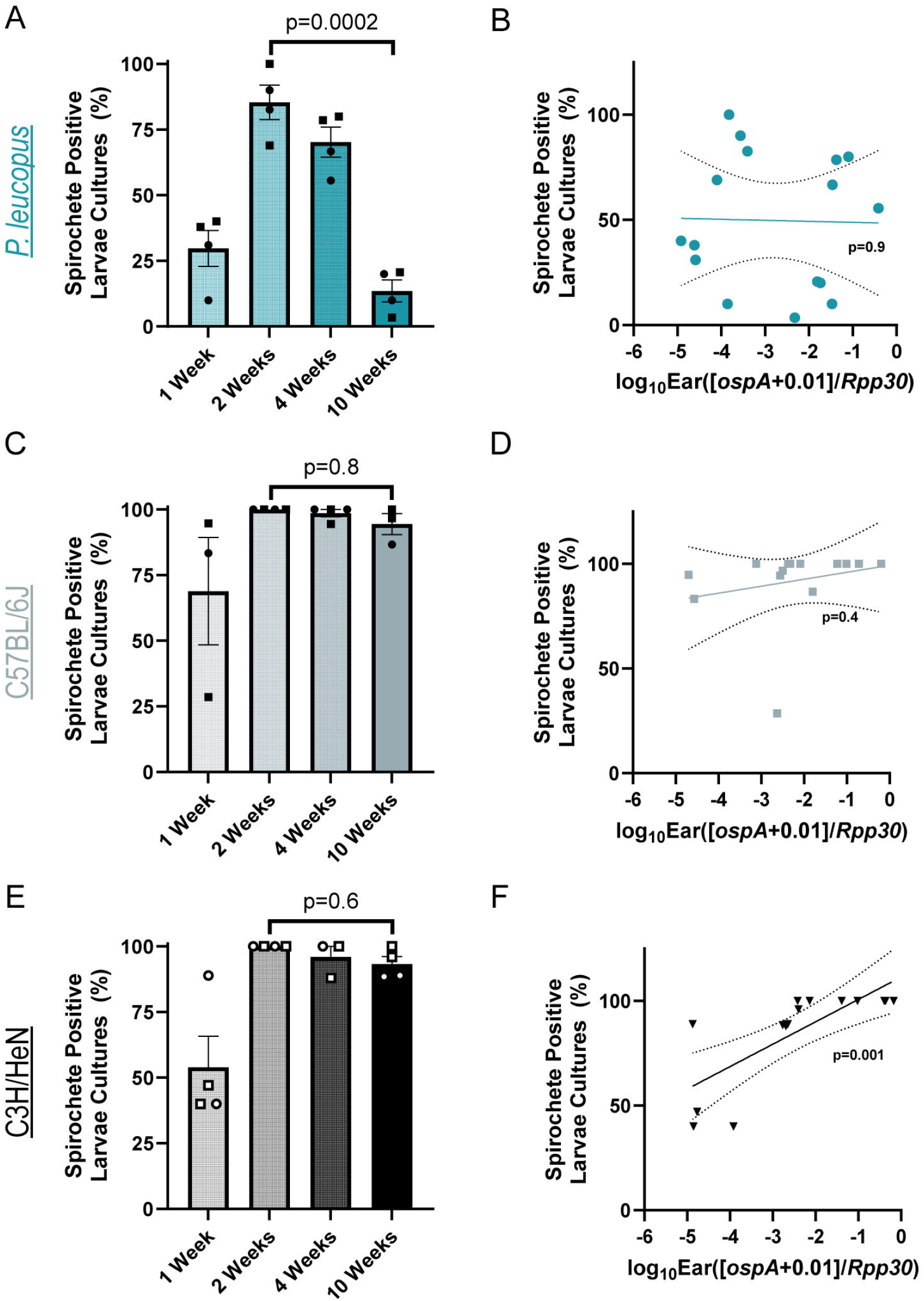
*P. leucopus* and *M. musculus B. burgdorferi* transmission to larval ticks over time. (A) *P. leucopus* have reduced transmission of *B. burgdorferi* to larval ticks over time. Rodents were infested with approximately 50 larval *I. scapularis* at 1-, 2-, 4-, and 10-weeks post-infestation with B31-infected nymphs. Fed larvae were crushed in BSK-II and monitored for growth to determine the rate of transmission from the rodents to *I. scapularis*. (B) In *P. leucopus, B. burgdorferi* burden in the ear (measured by ddPCR) does not correlate with transmission. Ear punches were taken just before infestation with larval ticks. (C, D) C57BL/6J mice remain highly transmissive over time (C) and burden in the ear (measured by ddPCR) does not correlate with transmission (D). (E, F) C3H/HeN mice remain highly transmissive over time (E) and burden in the ear (measured by ddPCR) does correlate with transmission (F). For panels A, C, D, the transmission was calculated as the number of fed larval ticks that yielded culturable spirochetes divided by the total number of fed larval ticks collected (5-30 fed larval ticks per rodent were collected, average=20.5 ticks/rodent). Rodents with fewer than 5 fed larval ticks collected were excluded. Each dot is an individual rodent (circles = female, squares = male) and bars mark the arithmetic mean. Reported p-value is from a one-way ANOVA on the log-transformed data with Šidák’s multiple comparison test comparing 2-weeks to 10-weeks to test for reduced infectivity over time. For panels B, D, F, each dot is an individual rodent, the error bars mark the 95% confidence interval of the slope, and the p-value is from a Pearson r correlation.

Examining rodent-to-tick transmission in laboratory mouse strains revealed a substantially different pattern. In C57BL/6J mice, ∼70% of larvae became infected when placed one week after nymph placement, 100% were infected when placed after two weeks, and >90% were infected when placed thereafter (two-week vs ten-week timepoint, p=0.8) **(Figure 3C**). Similar to *P. leucopus*, there was no relationship between the bacterial burden in the ear and rodent-to-tick transmission **(Figure 3D)**. Curiously, while C3H/HeN mice had very similar results to C57BL/6J mice in terms of transmission dynamics **(Figure 3E)**, we did actually observe a correlation between the burden in ear and transmission (**Figure 3F**, p=0.001, r=0.7586; Pearson Correlation). This aligns with a recent study which found a relationship between ear burden and rodent-to-tick transmission in C3H/HeJ mice (33). Regardless, these data collectively demonstrate that the inbred *M. musculus* strains used here transmit *B. burgdorferi* B31 to ticks more efficiently and over longer periods of time than *P. leucopus*.

### *P. leucopus* skin transcriptomics reveal a subtle but prolonged immune response against *B. burgdorferi*

Our previous results led us to hypothesize that *P. leucopus* has an efficient immune response that limits the bacterial burden in the skin and reduces transmission to ticks. To test this hypothesis, we performed RNA-seq on *P. leucopus* ear tissue from 1-, 2-, 4-, and 8-weeks post infestation. While bulk RNA-seq is a blunt tool which cannot always capture the complexities of an immune response, it is a useful technique to investigate *P. leucopus* immunity given the small number of validated molecular tools for this species.

Differential expression analysis identified a small number of upregulated and downregulated genes (p_adj_≤0.05) across time points **(Figure 4A-D, Supplemental File 1)**. This analysis revealed 4 differentially expressed genes at one week post infestation, 87 at two weeks, 24 at four weeks, and 20 at eight weeks **(Figure 4E)**. Most of these genes had increased expression in response to infection. Tracking these differentially expressed genes across the time course of the infection revealed that most genes upregulated at two weeks post infestation did not remain upregulated later during infection **(Figure 4F**). However, many of the genes that were still highly expressed at four weeks post infestation also had elevated expression at eight weeks post-infestation. Thus, it appears that after a burst of transcriptomic changes in skin of *P. leucopus* during acute infection, the transcriptomic response to *B. burgdorferi* B31 is refined and sustained into late infection.

**Figure 4:**
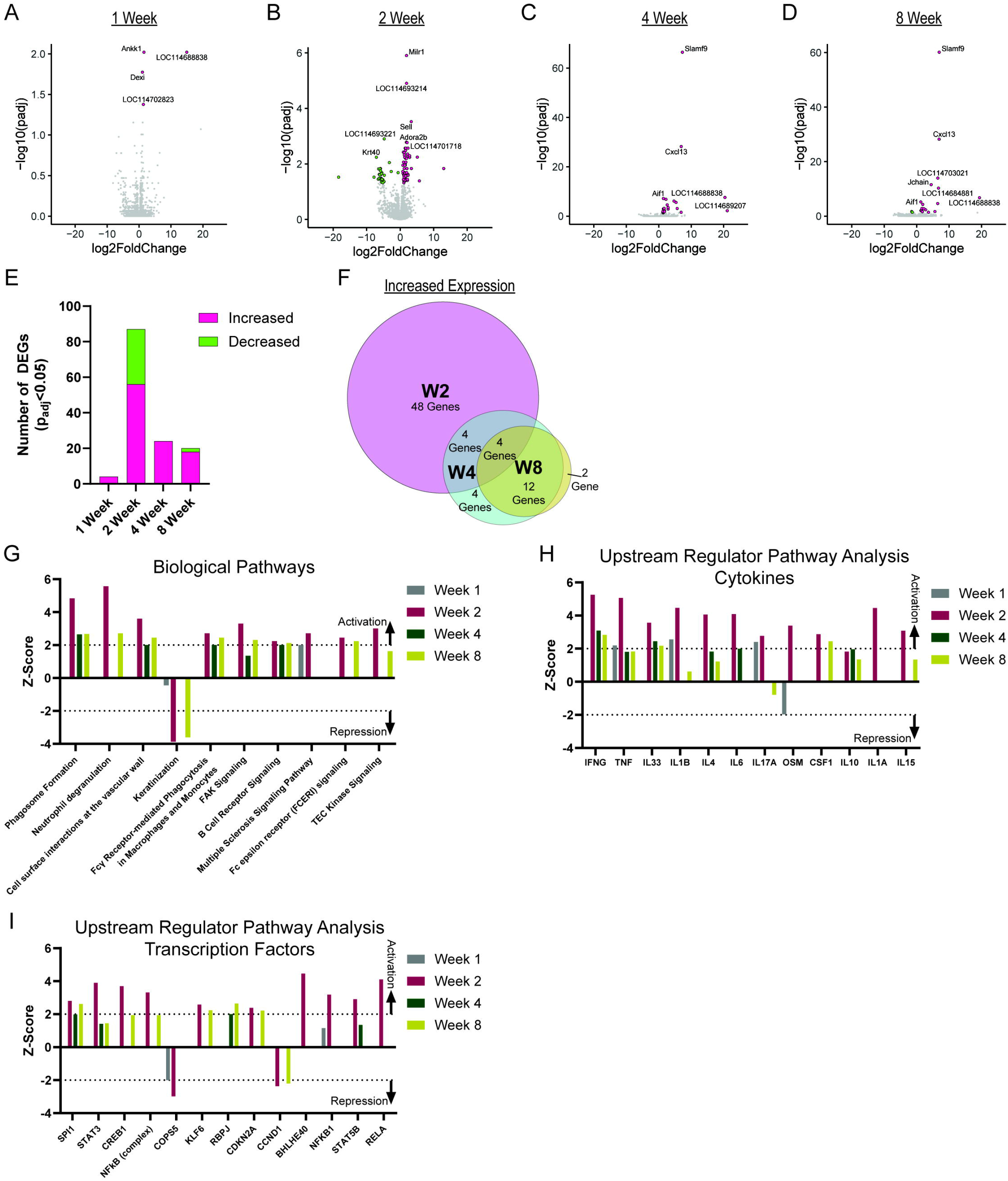
*P. leucopus* have a subtle but prolonged immune response to *B. burgdorferi* B31 in skin. (A-D) Differentially expressed genes in *P. leucopus* skin at 1- (A), 2- (B), 4- (C), and 8-weeks (D) post infestation compared to uninfected *P. leucopus*. Genes with p<0.05 are colored green or pink if down or up regulated, accordingly. (E) More genes are upregulated than downregulated across timepoints using a p<0.05 threshold. (F) Many upregulated genes overlap across timepoints. Overlap between gene sets was determined using the Biovenn software (54). (G-I) Use of QIAGEN IPA software (36) reveals activation or suppression of (G) biological pathways, (H) pathways regulated by cytokines, and (I) pathways regulated by transcription factors in *B. burgdorferi*-infected *P. leucopus*. For G-I, activation of pathways was determined using a Z score cutoff of 2 and repression was determined using a Z score cutoff of −2. RNA was derived from 4 *P. leucopus* rodents (two males, two females) at each timepoint.

We comment on a few differentially expressed genes. First, across two weeks, four weeks, and eight weeks post infestation, we observed upregulation of multiple C1q genes, which could lead to increased complement-mediated killing of *B. burgdorferi* at these time points. Additionally, we observed high expression of *Cxcl13* at four weeks and eight weeks post infestation, which suggests that resident cells are recruiting additional B cells to the skin to kill invading *B. burgdorferi*. Finally, we observed highly statistically significant (p_adj_≈10^-60^) increased expression of *Slamf9* at four weeks and eight weeks post infestation. Notably, this gene was also one of the top hits from Long *et al*.’s study on differential gene expression in the skin of *P. leucopus* at five weeks post intra-peritoneal *B. burgdorferi* infection (15). *Slamf9* has been associated with proinflammatory cytokine secretion in macrophages (34) and dendritic cells (35). This may suggest a role for these immune cells in mediating late-stage inflammation against *B. burgdorferi* B31 in skin.

We next utilized the Qiagen Ingenuity Pathway Analysis (IPA) pipeline (36) to understand which biological pathways are differentially regulated in *P. leucopus* in response to *B. burgdorferi* B31 infection. The IPA pipeline not only assigns a p-value for enrichment, but also attempts to leverage the direction of effect for each differentially expressed gene to determine whether the predicted pathway is turned on or off. We used this tool and identified enrichment of differentially expressed genes in multiple biological pathways associated with immunity that were activated at at least one timepoint **(Figure 4G)**, with some, including phagocytosis, macrophage activity, and B cell signaling showing evidence of activation at two weeks, four weeks, and eight weeks post infestation. Our analysis also identified activation of numerous cytokine-induced pathways at one or more timepoints, (including interferon-gamma (IFNG), tumor necrosis factor (TNF), oncostatin M (OSM), interleukin 1 beta (IL1β), and interleukin 6 (IL6)) **(Figure 4H**), as well as some immunity-associated transcription factor pathways (including SPI1, STAT3, NFκB, STAT5B, BHLHE40, KLF6, and RELA) **(Figure 4I)**. Note, this does not mean that the cytokine or transcription factor itself is differentially regulated in our dataset, but rather that genes known to be regulated by those upstream regulators are enriched in our dataset. Together, these data demonstrate that while we observe a small number of differentially expressed genes in the skin of *P. leucopus* following *B. burgdorferi* B31 infection, these genes provide a signature of an innate and adaptive immune response against the pathogen across timepoints.

### *P. leucopus* have very mild carditis following infection

Together, our data suggest that *P. leucopus* launch an immune response following infection, leading us to question whether these rodents display signs of inflammation at the tissue level. While *P. leucopus* do not have major behavioral (13, 14) or fitness (10, 14) changes in response to *B. burgdorferi* infection, with one dissenting study noting severe neurological symptoms that correlate with spirochete positivity (37), it remains possible that there are modest physiological changes in response to infection. Close examination of *P. leucopus* histology following infection is surprisingly sparse with some studies noting a lack of major inflammation in infected adults—though experimentally infected infant *P. leucopus* do develop arthritis (11). To examine whether *B. burgdorferi* induces an inflammatory response in infected tissues, we collected hearts from our laboratory-reared *P. leucopus* and the *M. musculus* strains at the terminal timepoints (**Figure 2A)** for histopathological analysis. Inflammation was assessed by a blinded pathologist that examined tissues for evidence of myocarditis, valvulitis, arteritis, and myocardial fibrosis.

In line with our observations that there is a modest but sustained inflammatory response in skin, we observed very mild carditis in *P. leucopus* four weeks post infestation **(Figure S1A, File S2)**. This inflammation was substantially lower than what we observed in C3H/HeN mice **(Figure S1B, File S2)**, and closer to what we observe with C57BL/6J mice **(Figure S1C, File S2)**. We note that the inflammation in *P. leucopus* is primarily characterized by myocarditis, with some rodents displaying moderate valvulitis and arteritis **(Fig S1D, File S2)**. This suggests that there is a mild but detectable inflammatory response against *B. burgdorferi* B31 in *P. leucopus* hearts.

## Discussion

The primary goals of this work were to understand (1) how *P. leucopus* respond to infection with *B. burgdorferi* B31, and (2) how this response compares to the traditional *M. musculus* lab mouse model of infection. Historically, the lab mouse has been used to model manifestations of human Lyme disease and as a representative model of mammalian-*B. burgdorferi*-tick infection dynamics. However, our data suggest that *M. musculus* are a poor model of *P. leucopus*-*B. burgdorferi* interactions, as *B. burgdorferi* B31 has dramatically different infection dynamics in the two systems and has reduced capacity to infect larval *I. scapularis* in laboratory-reared *P. leucopus*. While *M. musculus* and muroids can serve as competent hosts for *B. burgdorferi*—particularly in Europe—North American *B. burgdorferi* strains predominantly coexist with *P. leucopus* and shrews (8), and thus we were surprised to observe higher fitness of *B. burgdorferi* B31 (a North American strain) in the non-native hosts (*M. musculus*). We consider these results to be counter intuitive as we anticipated that the ability for *B. burgdorferi* to persist in nature would have depended on potent and sustained susceptibility of *P. leucopus* to *B. burgdorferi* infection across all strains—including B31.

We are not the first group to attempt to measure experimentally infected *P. leucopus* (17, 18, 38-41) or *M. musculus* (18, 33, 42) transmission of *B. burgdorferi* to *I. scapularis*. Indeed our measurements of rodent-to-larvae transmission align with prior observations using the JD1 strain of *B. burgdorferi* in that both studies find *P. leucopus* transmission peaks at two weeks post infection and becomes consistently worse thereafter (17), but curiously diverged substantially from a recent report which used the B31-5A4 clonal strain and found rodent-to-larvae transmission increased over the first five weeks of infection before slightly dropping eight weeks post infection (39). Notably our results did closely match with a previous study using B31 in *P. maniculatus* which found *P. leucopus* were less transmissive than C3H mice (29), as well as another study that used an OspC Type A strain (BL206) or a Type E strain (B348) in *P. leucopus* which also observed lower transmission than in C3H mice and reduced transmission over time (18). While the latter study did largely match our *P. leucopus* findings, they observed substantially worse rodent-to-larvae transmission in C3H mice over time than we did across similar timescales. This highlights that there is natural genetic diversity within OspC types that likely influences vertebrate to tick transmission. Supporting this, analysis of additional strains across the literature have found highly variable patterns in rodent (*P. leucopus* (38, 40, 41) or C3H *M. musculus* (33, 42)) to *I. scapularis* transmissibility over time.

Recent work by Zinck *et al*. has shown that there are substantial differences among *B. burgdorferi* strains in their ability to infect ticks from C3H/HeJ mice (33) and other work has demonstrated that there is strain specificity within different reservoir hosts (39, 43, 44). These data raise the possibility that some strains may be better or worse adapted to *P. leucopus*. Strain B31 in particular has been passaged in *Mus musculus* and may be specifically better adapted to *Mus* than *Peromyscus*. We do not expect that this is the case, based on alignment of our work with the studies highlighted above. However, even if our phenotypes are driven in part from this artificial selection, it would highlight differences in the pressures on *B. burgdorferi* survival and transmission in each species. Screening additional *B. burgdorferi* strains in *P. leucopus* could enable identification of variable spirochete loci that associate with higher transmission in this host. Understanding the impacts of those theoretical spirochete loci on transmission from other vertebrate hosts could broaden our understanding of *B. burgdorferi* evolution. It will also be interesting to examine whether genotypes that associate with enzootic spread align with loci that associate with disease severity in the dead-end human host (45).

In our RNA-sequencing dataset, similar to a past transcriptomic study (15), we note limited direct evidence of immunity (*e*.*g*. a lack of differentially expressed cytokines), which can likely be attributed to the relatively small number of infiltrating immune cells into the large numbers of epithelial and mesenchymal cells that make up the tissue. However, based on our data examining *B. burgdorferi* burden during the acute stages of *P. leucopus* infection, we hypothesize that the immune response we detect here contributes to limiting *B. burgdorferi* infection. While we observed overlap of increased *Slamf9* expression in *B. burgdorferi*-infected *P. leucopus*, this represents one of only a few similarities between our work and the study of Long *et al*. (15). Notably, a recent reanalysis of this dataset using the most recent *P. leucopus* genome build (citation pending, private communication with A. Barbour) revealed that *Cxcl13*, which was the second most differentially expressed gene in skin during late timepoints in this study **(Figure 4C, D)**, was also highly upregulated in the skin of infected *P. leucopus* in their study. We note numerous differences in the experimental design that may account for disparities between the studies: (1) differences in infection method (intra-peritoneal vs nymph), (2) differences in *B. burgdorferi* strain (Sh-2-82 vs B31), and (3) differences in *P. leucopus* source (LL Stock from the *Peromyscus* Genetic Stock Center vs the Tufts *Peromyscus* Colony). This third point highlights an interesting future direction—examining how natural diversity in *P. leucopus* impact *B. burgdorferi*-related phenotypes. Examining diverse populations of *P. leucopus* will not only allow us to assess how well our observations with the Tufts Colony rodents fully represent *P. leucopus* biology, but also potentially allow deployment of genome-wide screening approaches to better understand *P. leucopus* immunobiology.

We note that field studies have observed higher *Peromyscus* to *I. scapularis* transmission in nature (>90%) than we observe in this study (46). This may result from “immunosuppression” driven by environmental conditions (food stress, co-infections, pregnancy) that are absent in the laboratory (47, 48). Alternatively, reinfection of *P. leucopus* by subsequent nymphal feedings could contribute to a longer window of transmissibility to larval ticks. Regardless, the discrepancy between field studies and laboratory studies highlights a gap in knowledge in how *B. burgdorferi* persists in nature.

In sum, our data demonstrate that there are differences in the ability for *B. burgdorferi* to cycle between laboratory-reared *P. leucopus* or *M. musculus* and ticks. This highlights a tried-and-true principal of experimental design borrowed from the statistician George E. P. Box: “All models are wrong, but some are useful.” Here we propose that while *M. musculus* are a useful model for many aspects of *B. burgdorferi* infection, they may not accurately model the North American enzootic cycle. We hope that improving our understanding of how *P. leucopus* limit spread of *B. burgdorferi* B31 into new hosts could provide new insights into how we might break the cycle of infection in nature.

## Acknowledgements

The authors would like to thank Dr. Alan Barbour for numerous helpful conversations over the course of this project. RNA sequencing analyses were made possible by the Tufts High Performance Computing Cluster (https://it.tufts.edu/high-performance-computing). We thank past and present members of the Hu lab for helpful feedback during the formulation of this project and in assisting with the bioinformatic analyses, particularly Julie McCarthy and Peter Gwynne.

## Methods

### Bacterial Culture

*Borrelia burgdorferi* strain B31 (Hu Lab AD strain, routinely reisolated from sequential passages through C3H/HeN mice after obtaining the parental strain from Patricia A. Rosa) was cultured in Barbour-Stoenner-Kelley II (BSK-II) medium in stationary, sealed tubes at 32°C until mid-exponential phase growth (∼5×10^7^ spirochetes/mL). BSK-II was made according to existing protocols (Table 1, (49)) and quality controlled by confirming *B. burgdorferi* growth was unaffected across batches. Bacteria were monitored for growth, motility, and counted by pairing a Petroff-Hausser counter with darkfield microscopy. B31-AD stocks were passaged a maximum of one time *in vitro* following isolation from subcutaneously infected C3H/HeJ (Jackson Labs, Strain: 000659) skin two to four weeks post-infection in order to generate frozen stocks in 25% glycerol. Plasmid content was measured after thawing from frozen stocks. In order to limit environmental contamination when culturing spirochetes from rodent or tick tissue, phosphomycin disodium (100µg/mL, Sigma P5396), rifampicin (50µg/mL, Sigma R3501), and amphotericin B (5µg/mL, Sigma A4888) were added to the medium. Cultures from tissue were monitored weekly by darkfield microscopy and cultures were considered positive if spirochetes were observed within three weeks of culture start.

**Table 1:** BSK Reagent List.

| **Table 1: BSK Reagent List** |  |  |  |
| --- | --- | --- | --- |
| Product | Manufacturer | Product # | Amount (grams) per 2000mL Water |
| CMRL with L-glutamine | US Biologics | C5900 | 19.3 |
| Neopeptone | BD | 211681 | 11.1 |
| Yeastolate | Gibco | 255772 | 4.44 |
| Hepes acid | Fisher | BP310-100 | 13.32 |
| Dextrose | Fisher | BP350-500 | 11.1 |
| Sodium citrate | Fisher | S279-500 | 1.55 |
| Sodium pyruvate | Fisher | BP356-100 | 1.78 |
| N-acetyl-glucosamine | ThermoFisher | A13047.18 | 0.89 |
| Sodium bicarbonate | Fisher | BP328-500 | 4.88 |
| Gelatin B | Acros Organics | 61225-5000 | 1.06% Final Concentration (~400mL of a 7% autoclaved solution) |
| Rabbit serum (Heat Inactivated) | Gibco | 16120-099 | 5% Final Concentration (~131mL) |
| BSA | Millipore Sigma | 810037 | 5% Final Concentration |
| Sodium Hydroxide |  |  | To pH 7.6 |

### Rodent Maintenance, Use, and Ethical Statement

All animal procedures were approved by the Tufts University-Tufts Medical Center Institutional Animal Care and Use Committee (IACUC, Protocol #B2021-84). Euthanasia was performed in accordance with guidelines provided by the American Veterinary Medical Association (AVMA) and was approved by the Tufts University IACUC. Rodents were maintained by the Tufts Comparative Medicine Services. The *P. leucopus* colony was started by Dr. Sam Telford using wild captured rodents from the northeastern and midwestern United States. The colony has been closed since 1994, held in microisolator cages, and is specific pathogen free (regular sentinel testing).

With the exception of **Figure 1E**, all rodents were between 7 and 12 weeks old at the time of infection. Experiments were started with equal numbers of male and female rodents. Sample sizes are included in the figure legends for each panel. For **Figure 1E**, only male rodents were used and *P. leucopus* were >3 months and <10 months old, while both *M. musculus* strains were between 7 and 12 weeks old. No rodents were excluded from analysis unless either (1) unexpected rodent death or (2) failed nucleic acid extraction prevented data collection.

### Generation of *Ixodes scapularis* Nymphs

Immunocompromised C3H *scid* mice (C3SnSmn.Cg-Prkdcscid/J, Jackson Labs, Strain: 001131) were infected by subcutaneous injection of 1×10^5^ *B. burgdorferi* in 200 µL of BSK-II media. Two to eight weeks post-infection rodents were infested with larval *I. scapularis* (National Tick Research and Education Resource, Oklahoma State University, stored at 4°C in airtight containers containing saturated potassium sulfate). To infest rodents, ∼100 larval ticks were placed in modified 50 mL conical tube prior to placing a rodent in the tube. Rodents were restrained with ticks for one hour before returning to a cage suspended in a water moat. Water moats were monitored for ten days to collect fed larval ticks. Up to 6 fed larval ticks were stored in 1.7 mL microcentrifuge tubes with needle-generated airholes in an airtight glass desiccator containing water, allowed to molt, and stored for between 10 to 52 weeks post larval feeding. For each batch of nymphal ticks, six molted nymphs were crushed in BSK-II and monitored for spirochetes to confirm infection—and all six nymphs were consistently found to be infected. Uninfected nymphs were identically generated except for the use of uninfected C57BL/6J (Jackson Labs, Strain: 000664) mice and were confirmed to be uninfected by crushing in BSK-II.

### Rodent Infection with *B*. *burgdorferi*

*Mus musculus* (C57BL/6J [Jackson Labs, Strain: 000664], C3H/HeN [Charles River Laboratories, Strain: 025]) and *Peromyscus leucopus* (Sam Telford, Tufts University) were shaved prior to infection. The *P. leucopus* colony has been closed since 1994 and have been maintained under specific pathogen-free conditions. For needle infections, 200 µL of BSK-II containing 1×10^5^ bacteria were subcutaneously injected into each rodent at a dorsal posterior site. For nymph placements, a 1:4 mixture of melted beeswax to rosin gum mixture was heated and used to attach the top of a modified 1.7 mL microcentrifuge tube lined with mesh between the shoulder blades of the rodent. Five nymphal ticks were carefully placed through the mesh and into the placement cap. Four days later, caps were carefully removed, and ticks were collected. Rodents were singly housed in cages suspended in a water moat until all ticks were collected. Rodents were then returned to cohabitation with other infected animals of the same strain and sex.

### Determining Bacterial Burden

DNA from ears was collected by taking one to two 2-mm ear punches from one or both ears at listed time points. DNA from the tick bite or needle site was collected by dissection of a roughly 2-mm region in the center of the shaved region. In all cases, DNA was extracted using the Qiagen Blood and Tissue kit following overnight incubation at 56°C in Buffer ATL with Proteinase K. Following DNA extraction, digital droplet PCR (ddPCR) was performed to quantify the amount of *B. burgdorferi* DNA relative to rodent DNA. Primer probe pairs targeting mammalian *Rpp30* or *B. burgdorferi* are listed in **Table 2**. The primers targeting *Rpp30* are predicted to produce only a single product per gene copy per haplotype in both *Mus musculus* (Chromosome 19) and *Peromyscus leucopus* (Chromosome 1) (15, 50). Briefly, the Bio-Rad ddPCR Supermix for Probes (no dUTP) kit was used to perform the reaction within droplets generated by the QX200 droplet generator (Bio-Rad) in Oil-for-Probes (Bio-Rad). The reaction included 10 minutes at 95°C, 40 cycles of 94°C for 30 seconds and 60°C for 1 minute, followed by 10 minutes at 98°C and cooling to 4°C for at least 30 minutes to stabilize droplets. Fluorescence was calculated by the QX200 droplet reader and analyzed on the QX Manager Software (Bio-Rad).

**Table 2:**
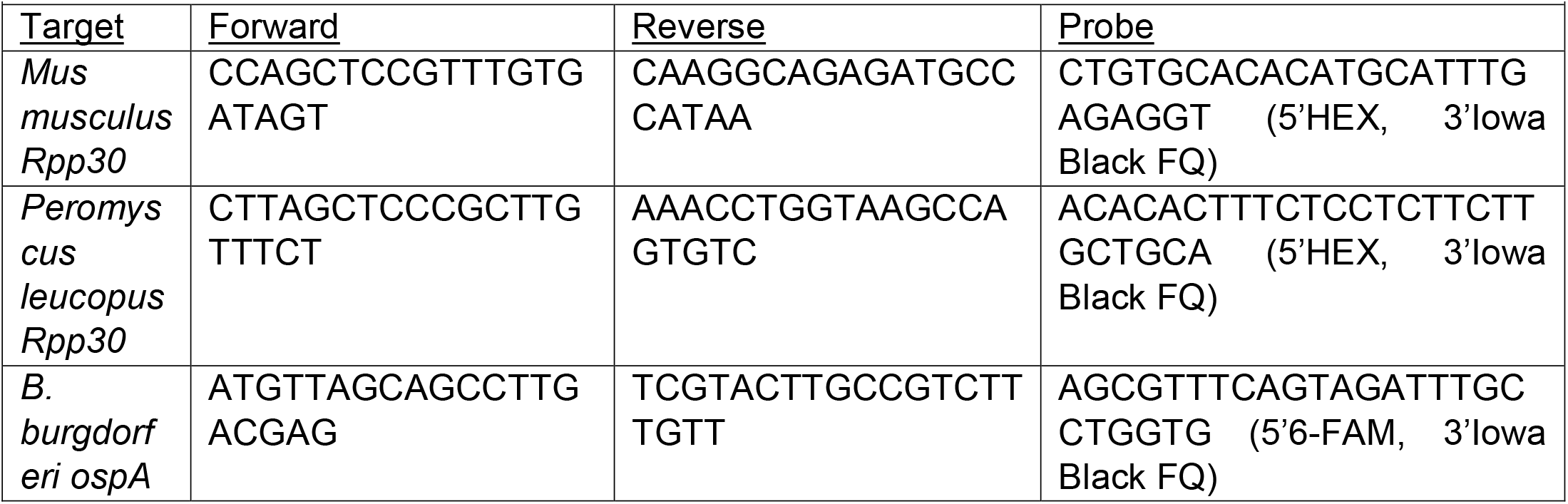
Oligonucleotides used in this study.

### RNA Extraction and Sequencing

One 2-mm ear punch from each ear was collected and submerged in RNA-later (Invitrogen) overnight at room temperature before being frozen at −80°C. After all samples were collected, samples were thawed on ice, transferred to QIAzol, and bead beaten with a 6.35 mm chrome steel bead (Biospec Products) for two cycles of two minutes each at an oscillation frequency of 30/second using a TissueLyser II (Qiagen). RNA was extracted using the miRNeasy mini kit (Qiagen), with the modification that two RWT washes were performed. gDNA was digested using TURBO DNase (Invitrogen) and RNA was repurified using the RNeasy MinElute Cleanup Kit (Qiagen). Purified RNA was submitted to Azenta Life Sciences/Genewiz for library preparation and sequencing (Illumina HiSeq 2×150bp). cDNA library was performed following polyA enrichment to deplete ribosomal RNA and we obtained approximately 20,000,000-40,000,000 paired-end reads per condition (median 25,593,087.5 paired-end reads, with a median 7,678 megabases sequenced).

### RNA Sequencing Analysis

Reads were aligned to the *P. leucopus* genome (GCF_004664715.2) (15) using STAR version 2.6.1 (51) and gene expression was summarized using RSEM version 1.3.1 (52). Annotations were based on the NCBI annotated genes. All differential gene expression analyses were performed with R (version 4.3.1) using DESeq2 version 1.40.2 (53) using standard parameters. Only genes with ≥10 reads across all conditions (summed) were included in the analysis. Differential gene expression was determined using DESeq2, modeling gene ∼ condition, where the condition term enabled comparison of transcripts from uninfected rodents against those in one of the timepoints (1 week, 2 weeks, 4 weeks, 8 weeks). The software leveraged a Wald test to generate p-values and a false discovery rate was computed using the Bejamini-Hochberg method. Genes with a false discovery rate ≤ 0.05 were considered significant. All differential gene expression data is included in **File S1**. Additional pathway analysis was performed with the use of QIAGEN IPA (QIAGEN Inc., https://digitalinsights.qiagen.com/IPA) (36) using genes with an adjusted p-value cutoff of <0.05.

### Determining Transmission from Rodents to Larval *Ixodes scapularis*

At each timepoint ∼50 larval ticks were placed in modified 50 mL conical tube prior to placing a rodent in the tube. Rodents were restrained with ticks for one hour before returning to a cage suspended in a water moat. Water moats were monitored at days three and four to collect fed larval ticks. Between 5 and 30 fed larvae were collected per rodent and stored in an airtight glass desiccator containing water for one week to enable *B. burgdorferi* replication in the bloodmeal. Ticks were submerged in 70% ethanol and each tick was crushed in 1 mL of BSK-II containing phosphomycin (100 µg/mL), rifampicin (50 µg/mL), and amphotericin B (5 µg/mL). Cultures were grown at 32°C, monitored weekly by darkfield microscopy, and cultures were considered positive if spirochetes were observed within three weeks of culture start.

### Histological Analysis

Whole hearts were fixed in Bouin’s fluid (Sigma-Aldrich), then bisected longitudinally and both halves paraffin embedded. Tissues were then sectioned and stained with hematoxylin and eosin (H&E; Cancer Diagnostics Inc) at the Comparative Pathology and Genomics Shared Resource at the Tufts Cummings School of Veterinary Medicine. Slides were blindly scored for myocarditis, valvulitis, arteritis and periarteritis, and myocardial fibrosis by a veterinary anatomic pathologist at Tufts as described below and an overall score (up to 10) was calculated for each animal. Specific definitions score criteria are defined as follows: **Myocarditis** [0 = no inflammation or occasional (≤3) clusters of 1-2 leukocytes (lymphocytes, histiocytes, or neutrophils) at the region of heart base (ventricular and atrial myocardium and epicardium; 1 = 1-2 clusters of 3+ leukocytes; 2 = 3-4 clusters of 3+ leukocytes; 3 = 5+ clusters of 3+ leukocytes], **Valvulitis** [0 = no inflammation or 1-2 leukocytes (lymphocytes, histiocytes, or neutrophils) within a valve (Valve leaflets and cusps (atrioventricular and semilunar)); 1 = 3-5 leukocytes; 2 = 6-10 leukocytes; 3 = 11+ leukocytes], **Arteritis and Periarteritis** [0 = no inflammation or occasional (≤3) clusters of 1-2 leukocytes (lymphocytes, histiocytes, or neutrophils) in the Aorta and pulmonary artery (tunica intima, media, or adventitia) and periarterial connective tissue; 1 = 1-2 clusters of 3-5 leukocytes; 2 = 3-4 clusters of 3-10 leukocytes or 1 cluster of 11+ leukocytes; 3 = 5+ clusters of 3-10 leukocytes or 2+ clusters of 11+ leukocytes], **Myocardial Fibrosis** [0 = absent in the region of heart base; 1 = present).

### Statistics

All statistical tests and sample size information for individual panels can be found in the figure legends. All statistical tests were performed in GraphPad Prism 10, with the exception of Figure 4 where tests were run either with R (version 4.3.1) using DESeq2 version 1.40.2 (53) or QIAGEN IPA (36). All ratios were log-transformed prior to use of Gaussian statistics (*e*.*g*. ANOVA). Post-hoc and/or multiple comparison testing was performed where appropriate, with the correction used for each panel listed in the figure legend. Where multiple comparison tested was used, corrected p-values or adjusted p-values are reported. All biological samples are from at least two independent experiments, defined as separate rodent infections performed on separate days.

### Data Availability

Sequencing data are deposited in GEO under accession number GSE244071.

---

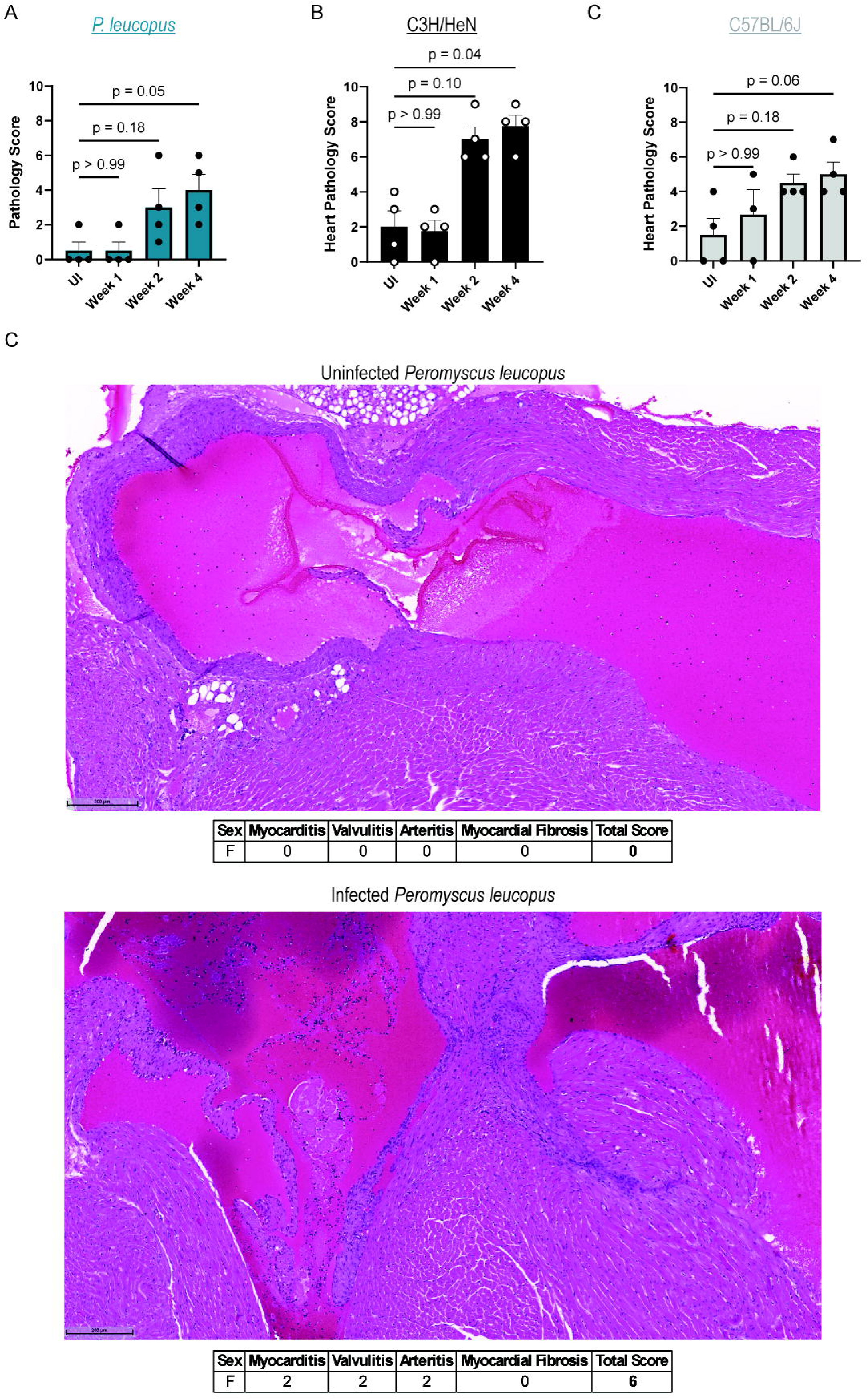

